# The third moments of the site frequency spectrum

**DOI:** 10.1101/109579

**Authors:** A. Klassmann, L. Ferretti

## Abstract

The analysis of patterns of segregating (i.e. polymorphic) sites in aligned sequences is routine in population genetics. Quantities of interest include the total number of segregating sites and the number of sites with mutations of different frequencies, the so-called *site frequency spectrum*. For neutrally evolving sequences, some classical results are available, including the expected value and variance of the spectrum in the Kingman coalescent model without recombination as calculated by Fu (1995).

In this work, we use similar techniques to compute the third moments of the site frequency spectrum without recombination. We also account for the linkage pattern of mutations, yielding the full haplotype spectrum of three polymorphic sites. Based on these results, we derive analytical results for the bias of Tajima’s *D* and other neutrality tests.

As an application, we obtain the second moments of the spectrum of linked sites, which is related to the neutral spectrum of chromosomal inversions and other structural variants. These moments can be used for the normalisation of new neutrality tests relying on these spectra.

## 1. Introduction

Statistics based on polymorphic loci are key to estimate relevant quantities in population genetics, such as the rescaled mutation rate *θ*. One common approach is to group variants together that appear with the same frequency in a sample and count the elements of each such group. The resulting summary statistic is called the *site frequency spectrum*.

The frequency spectrum is one of the most relevant statistics for population genetics. It can be used to infer evolutionary parameters such as mutation and recombination rate, past population history, demography and selection (Hud-son, 1983; Nielsen et al., 2005; Hein et al., 2004). Often, the variants are biallelic SNPs that can be “polarized”, e.g. for each site it is possible to say which allele is ancestral and which one is derived. This is the case for sequences with low mutation rate per base and for which an outgroup sequence is available. In what follows, we will consider exclusively this situation and assume that the evolution of these sequences can be modelled by a standard neutral Wright-Fisher model of constant population size.

Watterson (1975) credits Fisher (1930) with the first derivation (for a special case) of the first moments of the frequency spectrum. Their statement for the continuous case can be found in (Ewens, 1979), where it follows from results of di usion theory (Kimura, 1964). Watterson (1975) himself derived the first and second moments for the sum over all classes of the frequency spectrum, i.e. the number of segregating sites, using the technique of “moment estimators”. The full distribution of this quantity was shown by Tavaré (1984). The first and second moments for combinations of some components of the spectrum were later computed by Tajima (1989) using coalescent theory (Kingman, 1982) and combinatorics, while Fu (1995) completed this approach for the full frequency spectrum. A major application of his formulas is the normalisation of a class of neutrality tests such as Tajimas’s *D* (Tajima, 1989), as described by Achaz (2009). Recently, Hudson (2015) has given another proof of the first moments. As far as we know, higher moments of the spectrum have never been computed.

Asymptotic results for the distribution of the spectrum have been obtained by Dahmer and Kersting (2015). Their approach shows that for sample size *n* → ∞ and large *θ* (i.e. ignoring mutational Poisson noise), the first *k* components of the spectrum converge in distribution to i.i.d normal variables with mean *θ*/*i* and variance *θ*^2^ In (*n*)/*n*. However, their method applies only to allele counts which are much smaller than the sample size, hence it does not provide information on the full frequency spectrum in finite samples.

In this article we derive exact expressions for the third moments of the frequency spectrum. We use notation and approach of Fu (1995), with some technical modications in order to keep the number of different cases manageable. We derive independently by the approach of Watterson (1975) the third moment of the number of segregating sites and show the consistency of the two approaches. An immediate corollary of the third moments is the expected frequency spectrum for three linked segregating sites, which fully characterises the expected haplotype structure for triplets of sites.

We present two applications. The first one concerns the bias of neutrality tests. Several neutrality tests based on the frequency spectrum, like Tajima’s *D*, should ideally have an expected value of zero, yet they don’t. For the first time, we obtain general expressions for the bias of these neutrality tests as a function of mutation rate and sample size.

Finally, we derive the variance of the frequency spectra of nested and disjoint mutations at sites linked to a focal mutation. These spectra are equivalent to the spectrum of neutrally evolving chromosomal inversions (Ferretti et al., 2017). Moreover, they represent the basis for the derivation of the spectra of other structural variants. With these results, it is possible to obtain the proper normalisation for new Tajima’s *D*-like tests relying on the spectrum of linked mutations, e.g. neutrality tests for chromosomal introgressions or inversions.

## 2. Results

As is common practise in coalescent theory, we define *θ* as the population-scaled mutation rate per sequence, i.e.
*θ* = 2*pN_e_µL* where *p* is the ploidy, *N_e_* is the effective population size, *µ* is the mutation rate per generation per bp and *L* is the length of the sequence in base pairs. We refer to the number of mutations of size *i* in a sample of *n* sequences (i.e. the frequency spectrum) as *ξ_i_*.

The model that we consider is the Kingman coalescent, with an infinite-sites model of mutations. We assume no recombination, i.e. complete linkage among sites.

### 2.1. The third moments of the frequency spectrum

Our main result is an analytical expression for the third moments of the frequency spectrum.

**Theorem 1.**

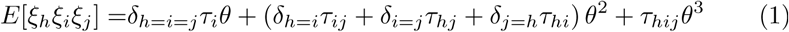

*for* 1≤ *h,i, j* < *n. The functions τ are:*

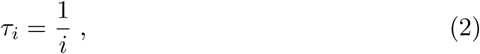

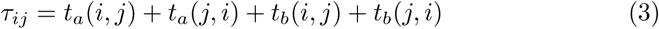

*with*

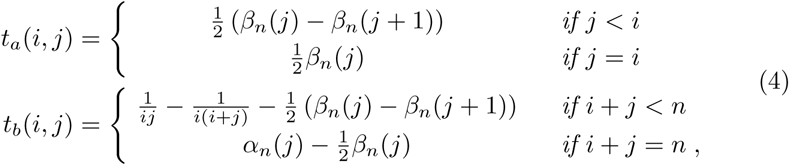

*and*^1^

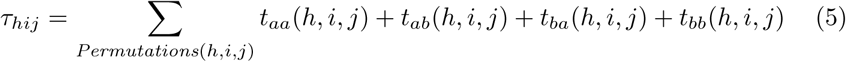

*with*

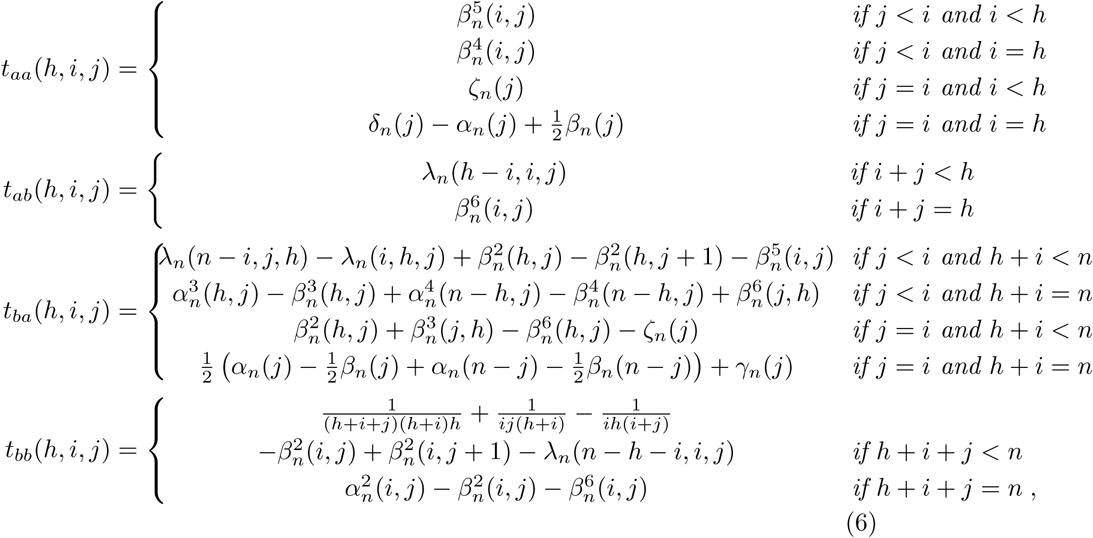

*using the following auxiliary functions (notation with upper indices):*

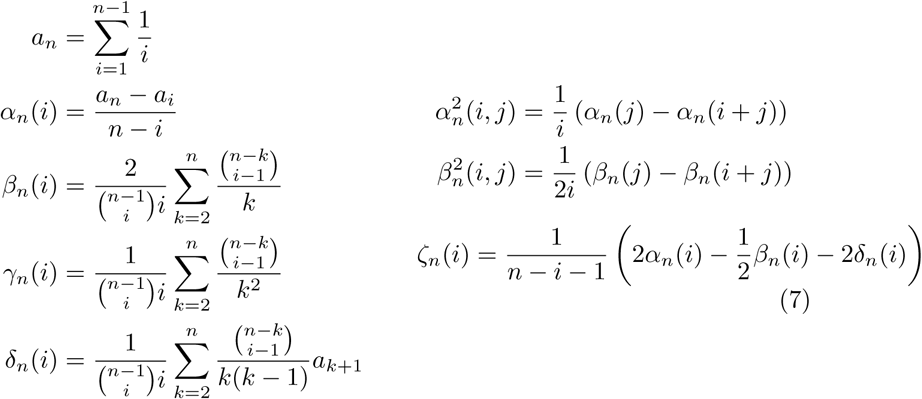

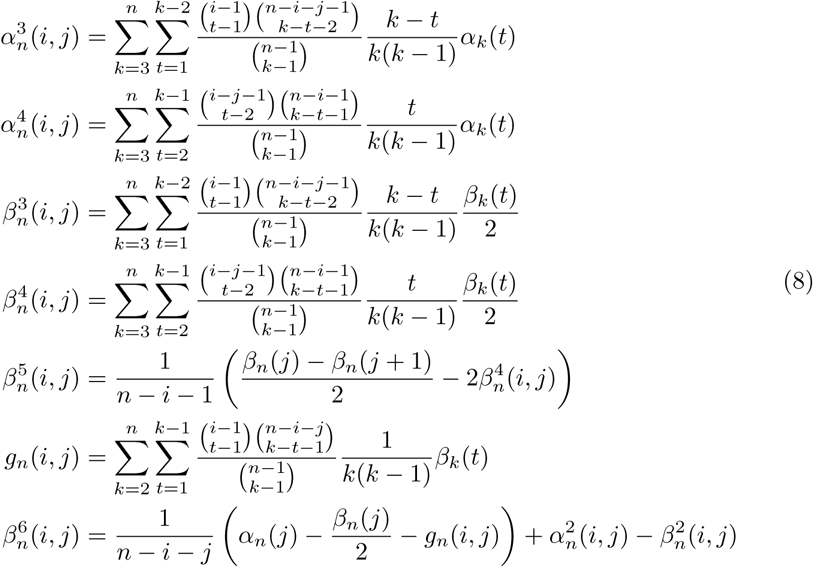

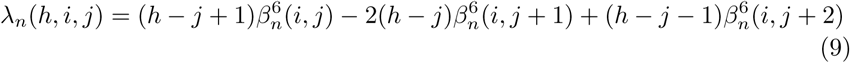

**Remark 1.** The coefficient for *θ* is the well known result for the expectation of the frequency spectrum

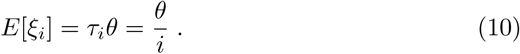

The terms *τ_ij_* are identical to the quadratic part of the second moments,

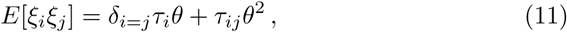

computed by Fu (1995): 
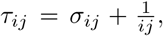
, with σ*_i,j_* defined in eq. (2) and (3) therein.

**Remark 2.** Fu (1995) showed in his eq. (34), that for *β_n_*(*i*) exists a more compact form, namely

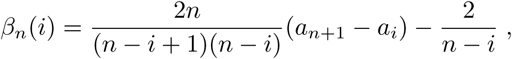

Where the summation over *k* is hidden in the *a_n_*. We do not have a similar form for the other expressions, however 
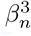
 and 
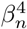
 can be expressed in terms of *g_n_*, too. Hence in a computational implementation the speed limiting factor are the double summations in 
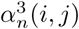
, 
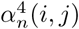
 and *g_n_*(*i,j*).

**Remark 3.** The sum over permutations simplifies the fractions in *t_b_* respective *t_bb_*

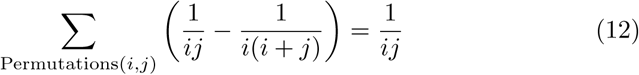

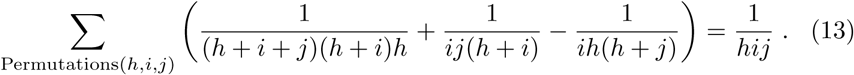

**Remark 4.** The central third moments can be obtained by

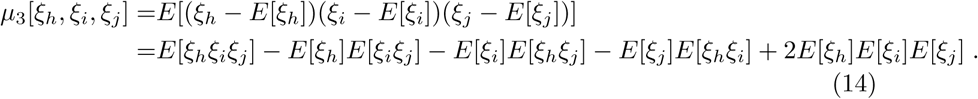

**Remark 5. For the folded spectrum**

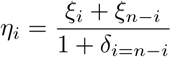

the corresponding third moments can be computed in a simnilar way as the second moments (eq. (9) in Fu (1995))

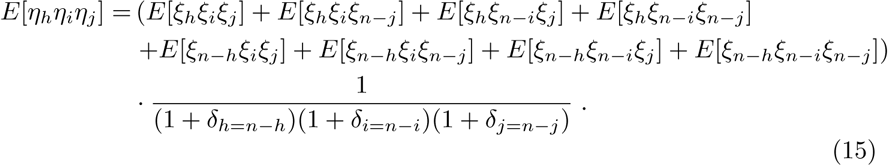

#### 2.2. The frequency spectrum of three linked sites

The components *t_aa_*, *t_ab_*, *t_ba_* and *t_bb_* correspond to different linkage patterns of three mutations (without recombination). The analogous pattern for two sites has been recently studied in Ferretti et al. (2017). Pairs of linked mutations are *nested*, when one mutation is present only in sequences containing the other, or *disjoint*, if the mutations are present in different sets of sequences. The nested and disjoint components of the frequency spectrum for pairs of sites give a complete description of the haplotype structure of two sites (up to permutations of individuals and sites).

Following the derivation in Ferretti et al. (2017), the frequency spectrum for triplets of segregating sites is given by

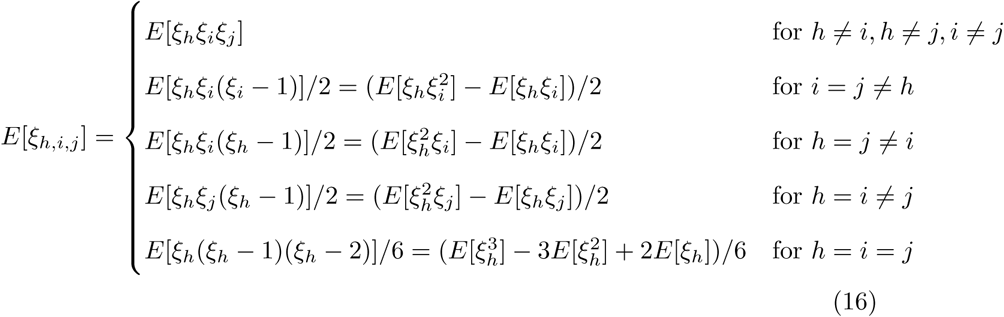

noticing that three derived mutations in different sites can have four possible relations:

- *fully nested*: the second mutation *i* is nested inside the first *h*, and the third *j* is nested inside the second *i*. This component corresponds to *t_aa_(h, i, j)*.
- *disjoint within nested:* the second and third mutations *i*, *j* are disjoint, but both are nested inside the first *h*. This component corresponds to *t_ab_(h, i, j)*.
- *nested within disjoint:* the mutations *h*, *i* are mutually disjoint, but mutation *j* is nested inside *i* (and consequently *j* is disjoint to *h*, too). This component corresponds to *t_ba_(h, i, j)*.
- *fully disjoint:* all mutations *h, i, j* are mutually disjoint. This component corresponds to *t_bb_(h, i, j)*.

Therefore, the spectrum of three sites can be easily decomposed by separating the components *t_aa_*, *t_ab_*, *t_ba_* and *t_bb_*. This spectrum is equivalent to a complete characterization of the haplotype spectrum of three sites.

#### 2.3. Comparison with the third moments of the number of segregating sites

We derive by the method of Watterson (1975) the third moments for the number of segregating sites 
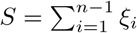

**Theorem 2.** *Writing* 
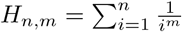
 *for the n-th harmonic number of order m, the third moment (resp. central moment) of the number of segregating sites S for a sample of size n is:*

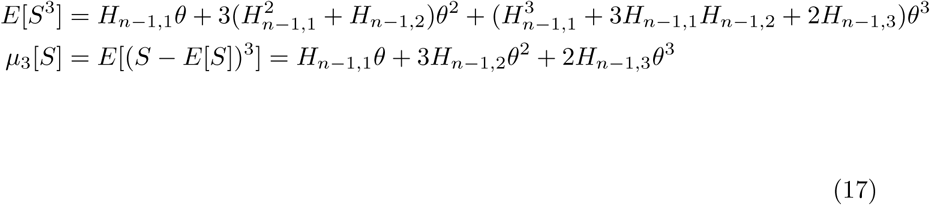

Since

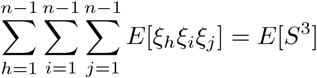

and

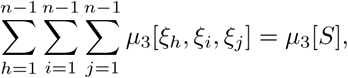

following from Theorem 1 and Theorem 2, the corresponding coefficients for *θ*, *θ*^2^ and *θ*^3^ have to be the same. We give in the supplement an explicit proof of the non-trivial identities of the coe cients for *θ*^2^ and *θ*^3^ stated as:
**Lemma 1**.

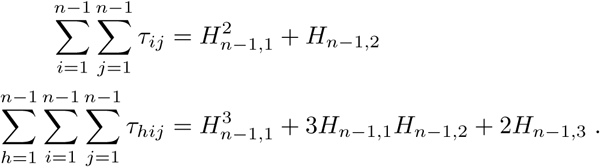

### 2.4. Skewness and bias of Tajima’s D and similar neutrality tests

One of the applications of the frequency spectrum is to test if the observed patterns in sequences are compatible with neutral evolutionary models. Several neutrality tests fall into a general class that relies on normalized linear combinations of the frequency spectrum (Achaz (2009), Ferretti et al. (2010)). Their general form is

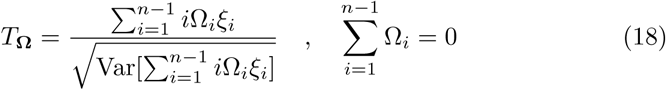

where the variance in the denominator

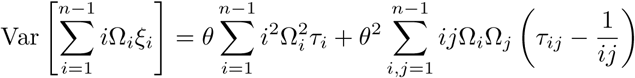

is a linear combination of *θ* and *θ*^2^. These two quantities, if unknown, are usually estimated from *S* and *S*^2^ by the method of moments: 
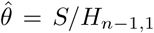
 and 
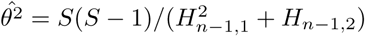
.

In this section, we explore the additional information that the third moments of the spectrum reveal about the distribution of neutrality tests, in particular about their skewness and bias.

**Table 1.** Weights and references of the analysed neutrality tests.

| Test | weights $\Omega_i$ | reference |
| --- | --- | --- |
| $D_{(Tajima)}$ | $(n-i)/\binom{n}{2} - 1/ia_n$ | TAJIMA (1989) |
| $D_{(Fu\&Li)}$ | $1/ia_n - \delta_{i,1}$ | FU and LI (1993) |
| $F_{(Fu\&Li)}$ | $(n-i) - \delta_{i,1}$ | FU and LI (1993) |
| $H_{(Fay\&Wu)}$ | $(n-2i)/\binom{n}{2}$ | FAY and WU (2000) |
| $E_{(Zeng)}$ | $1/(n-1) - 1/ia_n$ | ZENG *et al.* (2006) |

First, we consider the case of known *θ*. It is well known the distributions of neutrality tests based on the frequency spectrum such as Tajima’s *D* (Tajima, 1989) tend to be skewed (Hudson, 1991). These tests are normalized to mean 0 and variance 1 under the neutral coalescent with constant population size: *E*[*T*_Ω_] = 0 and Var[*T*_Ω_] = 1. Consequently, the skewness γ = *µ_3_*/ σ^3^ equals the third moment of the test:

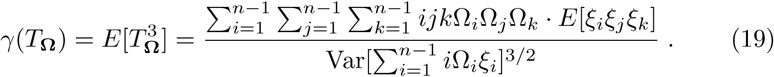

The weights Ω_*i*_ for some commonly used neutrality tests are given in Table 1. Figure 1 shows, that analytical results and those from simulations with ‘ms’ (Hudson, 2002) agree well. However, when the parameter *θ* has to be estimated from the data, as it is usually the case, the denominator of the test is a function of the estimator, contributing to the skewness. This has a relatively large effect, but surprisingly for most considered values of *θ* it reduces the skewness.

**Figure 1:**
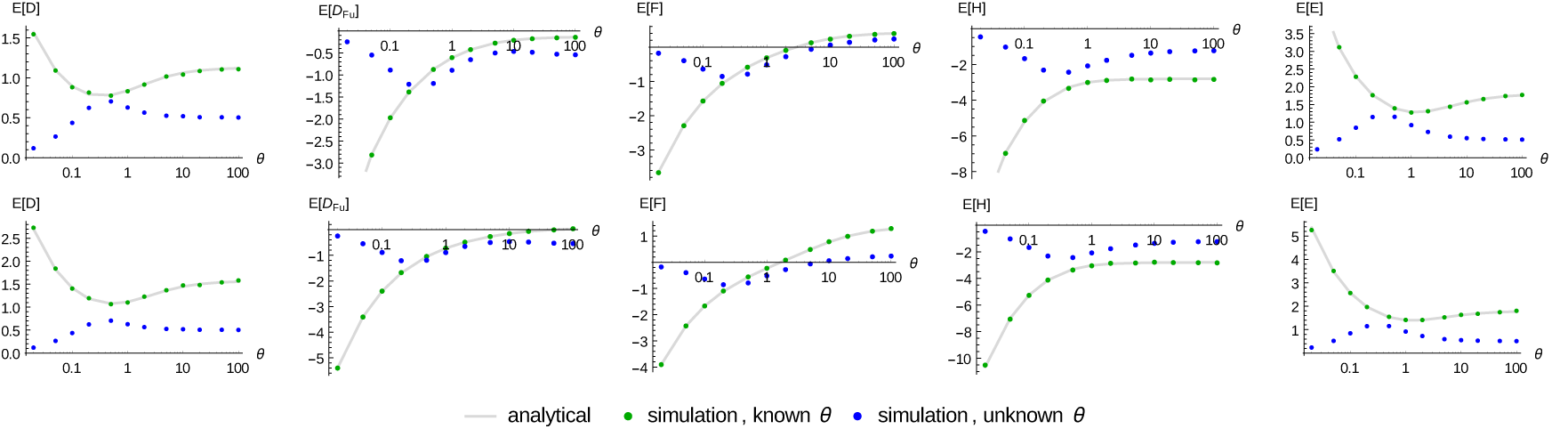
Skewness of neutrality tests for sample size *n* = 50 (top) and *n* = 500 (bottom). The analytical skewness was obtained by Eq. 19. For simulations, the skewness was estimated by 
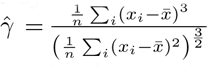
 over 10^6^ genealogies. The test values were calculated using the true *θ* (green points) and Wattersons estimator 
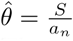
 (blue points), respectively.

**Figure 2:**
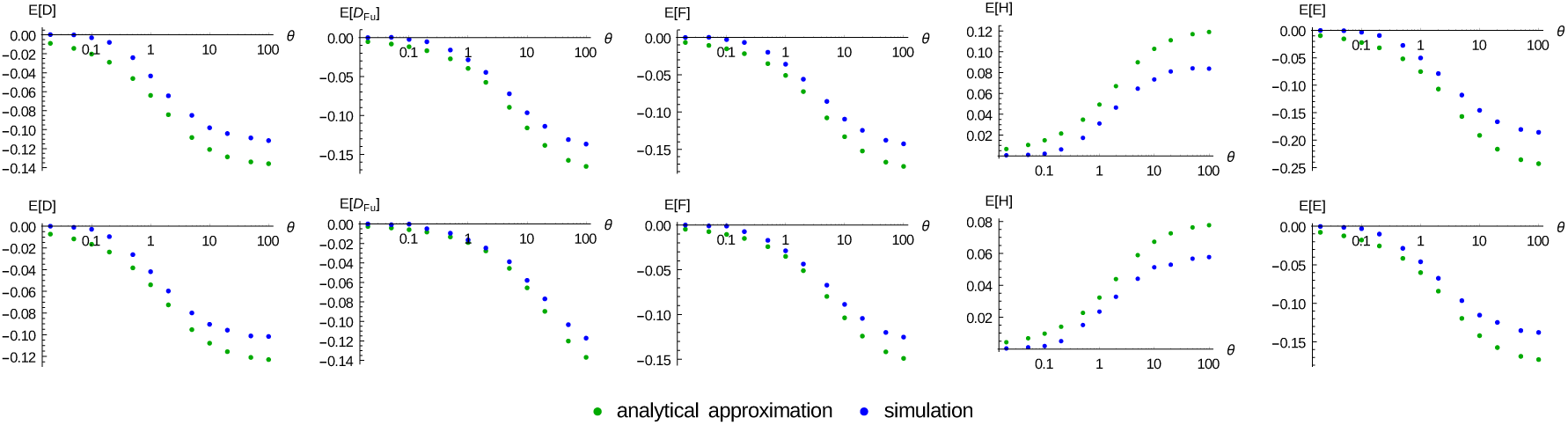
The bias of the tests mentioned in table 1 with sample size *n* = 50 (top) and *n* = 500 (bottom). Shown are the values of our analytical approximation and numerical data, obtained by simulation with’ms’, averaged over 10^6^ genealogies.

For *θ* unknown and estimated from *S*, we can still make use of the third moments. In this case, we can compute an approximate result for the bias of the test. We apply the following formula for the Taylor expansion of moments of random variables^2^ *X*, *Y* with *E*[*X*] = 0 and *Y* > 0

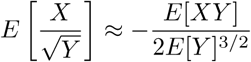

and the fact that 
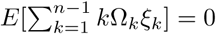
to obtain the bias:

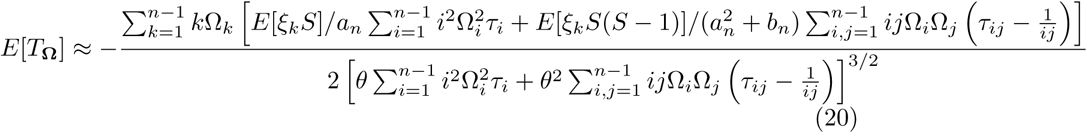

with 
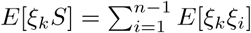
 resp.
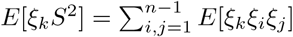
.

The above equation gives a reasonably good estimate of the bias of neutrality tests (Figure 2), taking into account that eq. (20) represents only the first term of a bivariate Taylor expansion.

### 2.5. The variance of the frequency spectrum of linked sites

We will use the nomenclature introduced by Sargsyan (2015) and expanded in Ferretti et al. (2017). We call a certain mutation of interest *focal* and we refer to it as *φ*. As above, further mutations that appear in at least one individual together with it, are called *nested* while all others are called *disjoint*. Note, that the focal mutation *φ* itself is not included into neither. More specically, we refer to the number of mutations of size *i* that are nested with the focal mutation by 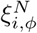
 and to those that are disjoint by 
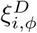
. Evidently, the number of overall occurrences of mutations of size *i*, given, *ø*, is 
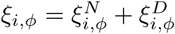
.

We now condition on the focal mutation *ø* being a mutation *ø* being a mutation of any size *h* and write: 
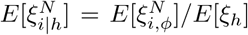
 resp. 
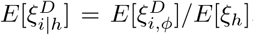
. The expectation values of nested mutations are a corollary of the second moments derived by Fu (1995); they are given in Ferretti et al. (2017). The second moments of two nested respective two disjoint mutations, or between one nested and one disjoint mutation, are obtained directly from the above expressions of the third moments:

**Corollary 1.**

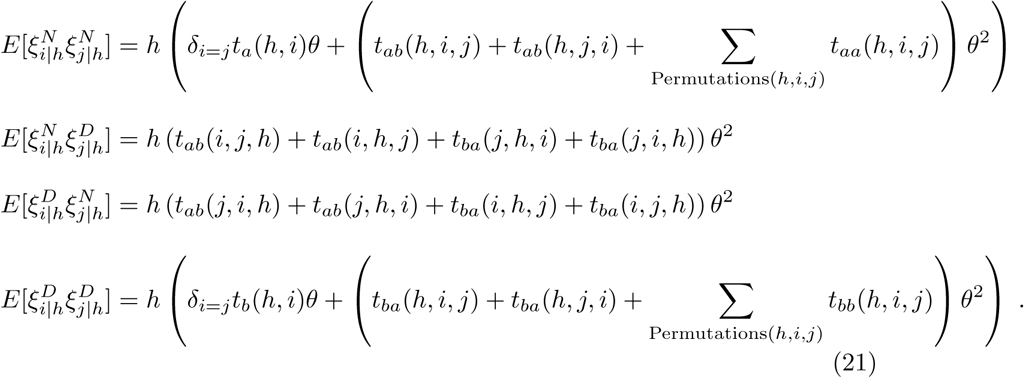

### 2.6. Numerical results

In Figure 4 we compare the analytical results with the third moments from coalescent simulations. We use “ms” (Hudson, 2002) to generate samples and from their frequency spectra we calculate estimates of the third moments. For increasing sample size *n* the “off-diagonal” elements of the three-dimensional array of third moments get increasingly small; that’s why the maximum relative error of the simulated data increases with *n*. The graphs show that with increasing number of samples, the values from simulations converge to our analytical results. However the convergence is extremely slow, indicating a large variance of the third moments.

**Figure 3:**
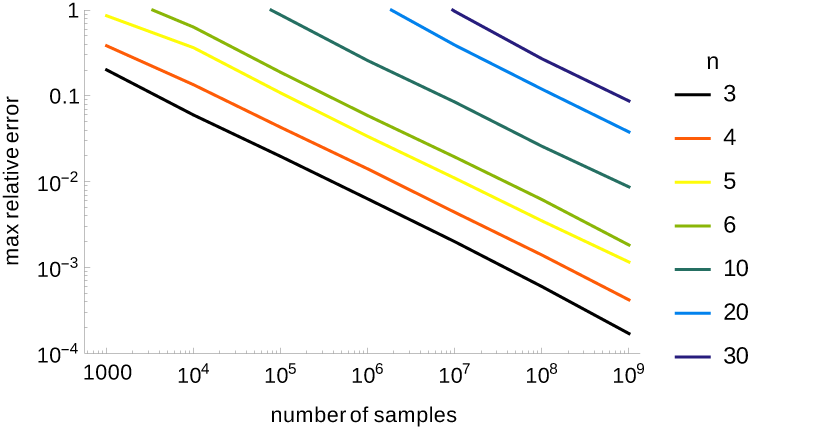
The relative error between expected (analytical) and observed (estimated from simulated data) third moments. We computed errors 
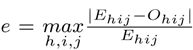
 where *O_hij_* was averaged over 10^3^ til 10^9^ genealogies (number of samples). The figure shows the average over 100 of these errors. The colours indicate different sample sizes *n*.

Figure 4 shows all third moments for a sample of size *n* = 5. As in the two-dimensional case, the values of the diagonals (now in 3 dimensions) dominate.

In Figure 5 we compare the covariances of the standard frequency spectrum with the covariances between nested and disjoint mutations of a conditional spectrum. The spectra of nested, resp. disjoint, sites are still dominated by the variances, while the correlation of “mirror sites” (*ξi* and *ξn–i* in the standard spectrum), is lost. There is almost no correlation between nested and disjoint sites.

**Figure 4:**
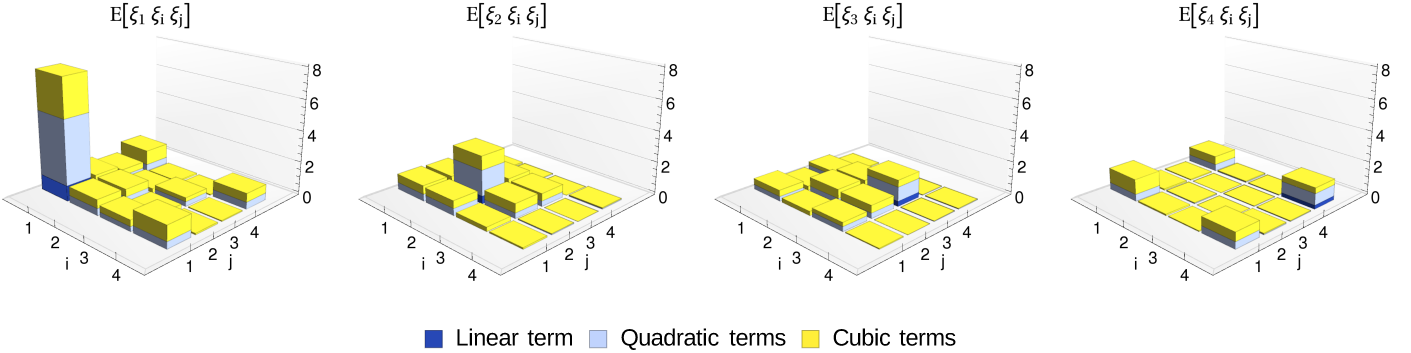
The expected values of all third moments for *n* = 5, *θ* = 1 and the respective contributions of the linear, quadratic and cubic terms.

**Figure 5:**
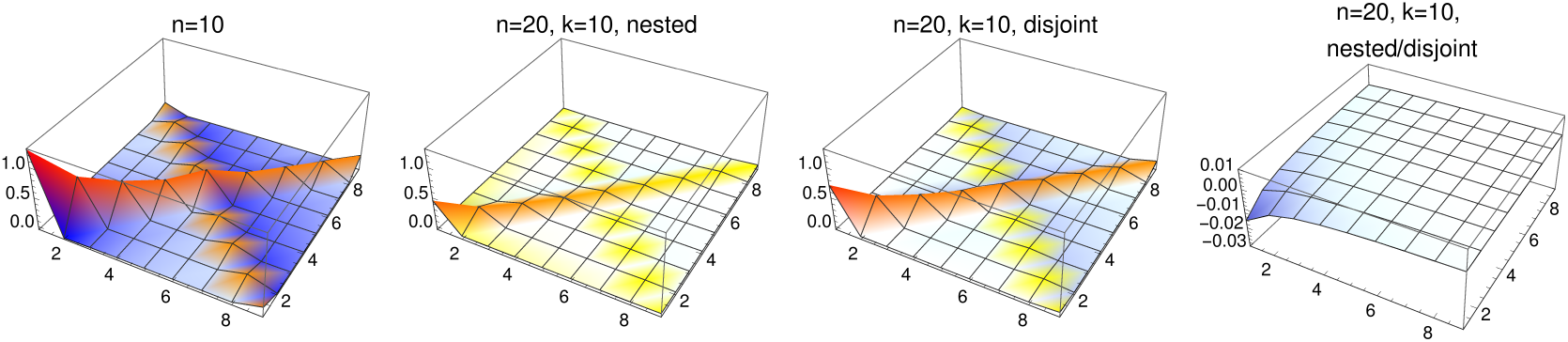
Comparison between unconditional and conditional covariances. Left: unconditional covariances *Cov*[*ξ_i_*, *ξ_j_*] for sample size *n* = 10. The remainder graphs show the covariances between mutations conditional on a mutation of size *k* = 10 in a sample of size *n* = 20. The left middle shows the covariances between mutations nested within the focal mutation, the right middle the covariances of mutations both disjoint and the rightmost the covariance between nested and disjoint mutations.

### 2.7. Comparison with asymptotic analytical results

Dahmer and Kersting (2015) showed the convergence of the distribution of the components of the spectrum to centered and rescaled i.i.d. Gaussian variables in the large *n* limit. More precisely, they state that for large, *θ* i.e. ignoring the Poisson noise, we have for fixed *k*

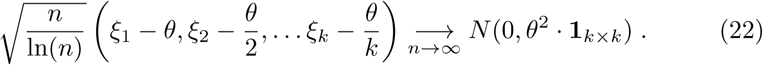

One could naively assume, that this means that in the limit of large *n* the *ξ_k_*s could be treated as independent Gaussian random variables with mean *θ*/*k* and variance *θ*^2^ In(*n*)/*n*, leading to the approximation

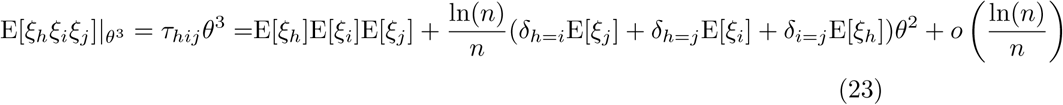

This is however incorrect. The distribution of each component of the spectrum *ξ_k_* shows excesses of outliers and heavy tails (Janson and Kersting, 2011), hence the convergence in distribution proved by Dahmer and Kersting does not imply the scaling of the moments.

Figure 6 shows, that the asymptotics are of limited help for a particular finite sample size *n*, since only moments for 
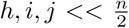
, and only those with at least two indices differing, seem to be approximated reasonably well.

**Figure 6:**
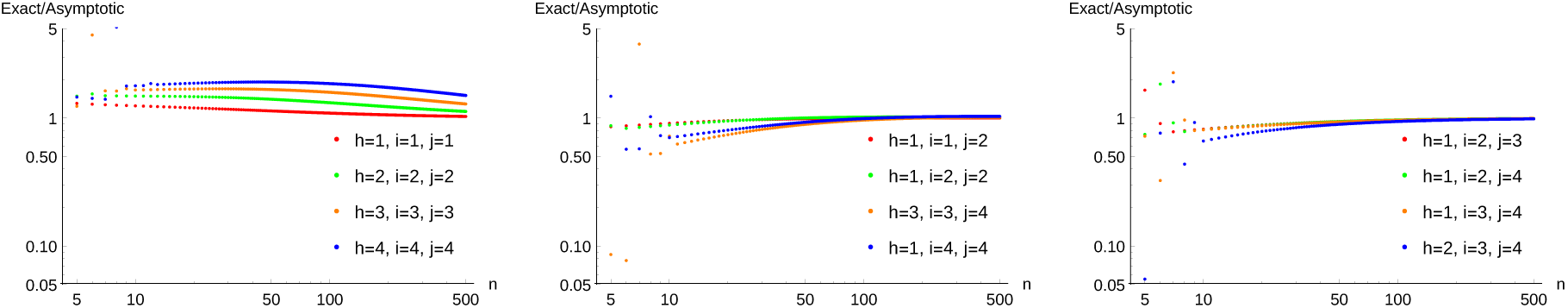
From (Dahmer and Kersting, 2015) follows, that mutations of small size within a large sample are approximately independent. Shown is the ratio of our exact results (*τ_hij_θ*^3^) to the asymptotic approximation (eq. (23)) for small fixed indices 1 ≤ *h*, *i*, *j* ≤ 4 and varying sample size *n*. Left: all indices are the same; middle: two indices differ; right: all indices differ.

## 3. Methods

### 3.1. Proof of theorem 1

#### 3.1.1. Separation of estimation

A coalescent tree is constructed by two independent stochastic processes, namely its branching pattern (the topology) and the lengths of its branches (coalescent times). The idea of Fu (1995) is to decompose the tree into small parts, called *lines*, by cutting each branch along *states*, which are delineated by coalescent events. He first calculates all possible hierarchical relationships between those lines, thereby transforming a probabilistic problem into a combinatorial one. Second, he computes the estimated mutations on each line. This number is correlated between lines of the same state because of their shared lengths. The combined sum over the two quantities yields the desired second moments. We re-use method and notation of Fu (1995) with appropriate extensions. A thorough explanation of the main ingredients of his proof, albeit with somewhat different notation, has been given in Durrett (2008). An extended “reprint” of the more technical parts can be found in the supplement of our companion paper Ferretti et al. (2017).

**Figure 7:**
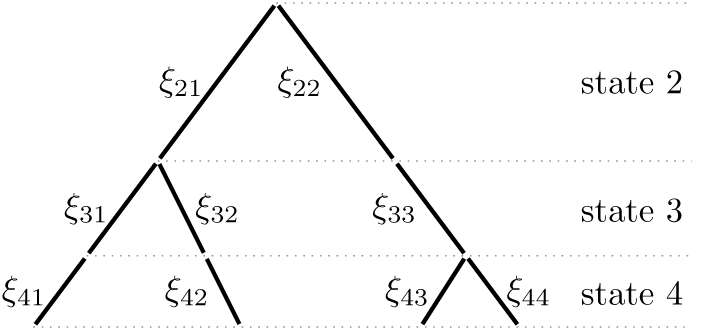
How do we calculate the expected number of mutations of size 2? For a tree with this topology *τ* only the lines *ξ*_21_, *ξ*_22_ and *ξ*_33_ have two descendants in the sample. Thus we have *∊*_21_ (2) = *∊*_22_ (2) = *∊*_33_ (1) = 1 and all other *∊_kl_*(2) are zero. It follows, that 
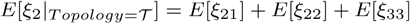
. *E*[*ξ_kl_*] is the expected amount of mutations on the line *ξ_kl_* which is proportional to its length. Averaging over all topologies yields *E*[*ξ_kl_*].

We define index variables *∊_kl_*(*i*) that indicate if the line *l* of state *k* has *i* descendants at state *n*, (e.g. they take the values 1 resp. 0). It follows that (cf. figure 7)

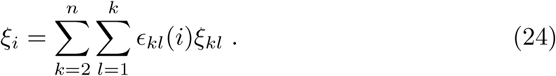

In the following we use the fact, that the index *l* serves only to distinguish lines of the same state, but otherwise has no meaning, since all lines of the same state are equivalent. The indicator variables are idempotent (*∊_kl_*(*i*)^2^ = *∊_kl_*(*i*)) and independent of the length (resp. mutation rate) *ξ_kl_*. The expectation values of the indicator variables correspond to probabilities, which we will define in the following subsection.

#### 3.1.2. Averaging over topologies

Figure 8: The possible hierachical relationships between three lines of a coalescent tree and their corresponding probabilities.

We split the computation of the expectation values of the indicator variables (which define the topology) into several cases, pictured in figure 8.

**Figure 8:**
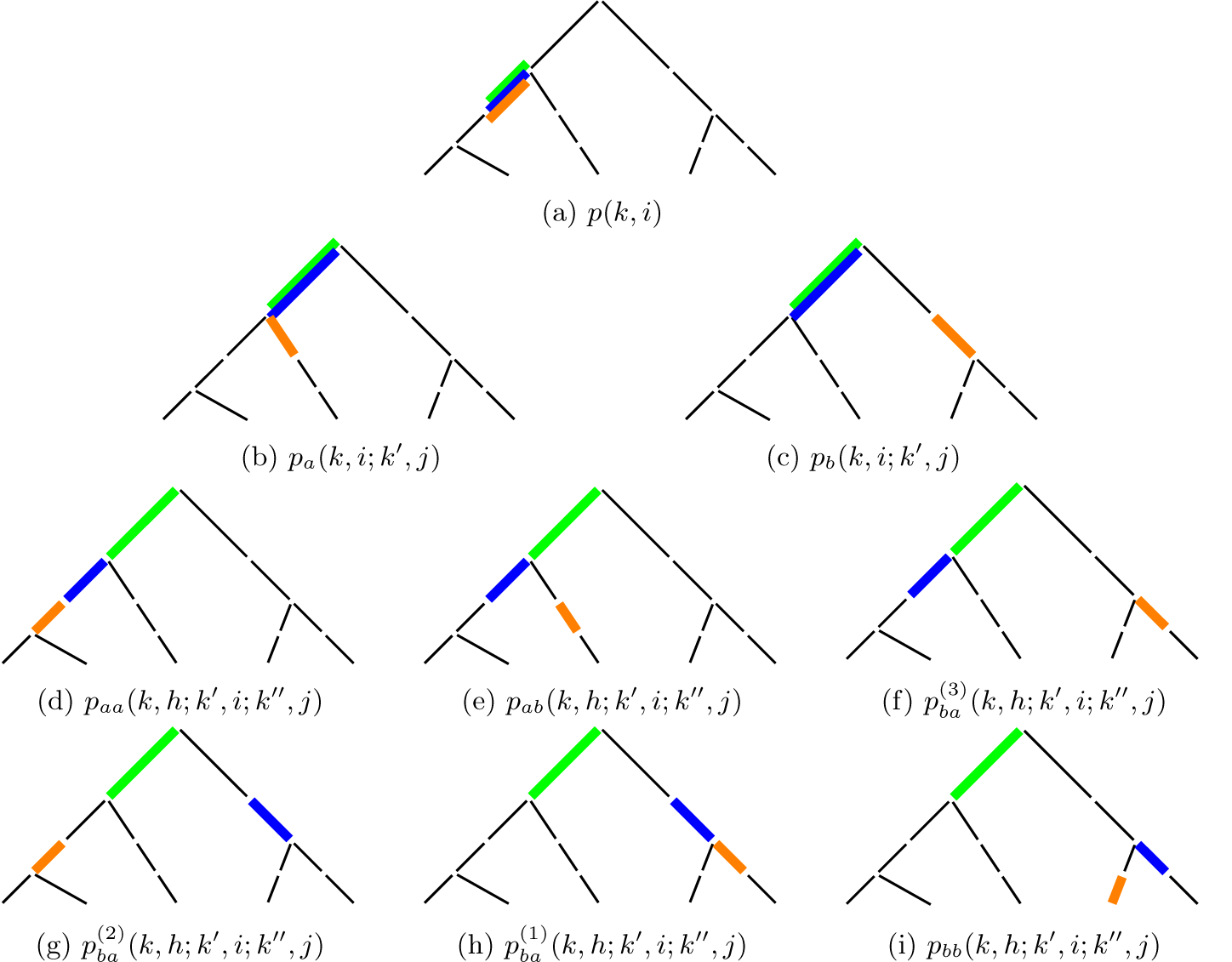
The possible hierachical relationships between three lines of a coalescent tree and
their corresponding probabilities.

We recall, that the number of descendants of lines in the coalescent is equivalent to that of balls of a specific colour in a so-called *Pólya urn model* whose probability distribution is known and reviewed in e.g. Griffiths and Tavare (2003). We introduce the following notation: *p*_*k* → *n*_(*t* → *i*) is the probability that *t* lines at state *k* have *i* descendants at state *n*. This probability is

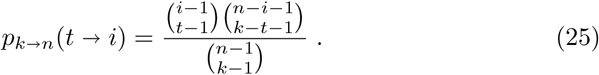

At this point it is helpful to define 
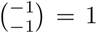
, while binomial coeffcients con-taining any other combination of one or two negative numbers are set to zero (Durrett, 2008). This makes it possible to subsume in the above and following formulas the case that *t* = *k* lines of state *k* yield *i* = *n* lines at state *n* (which is true with probability 1). Later on, these special cases will be resolved separately and none of the expressions in *Results* rely on this definition.

The probability that *t* and *u* (different) lines at state *k* have respectively *i* and *j* descendants at state *n* is

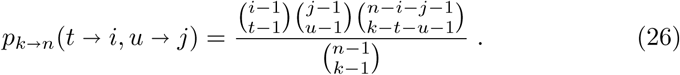

And for three such (non-overlapping) sets of lines the probability yields

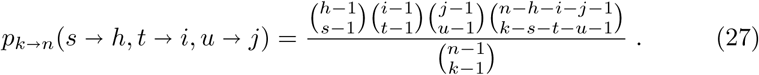

Using this notation we can now state the probabilities for different configurations. We start with those derived by Fu: The probability, that one line at state *k* has *i* descendants at state *n* is (Fu, 1995, eq. (14))

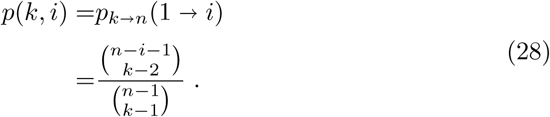

The joint probability that one line at state *k* and one nested line at state *k^′^* ≥ *k* have *i* respective *j* descendants at state *n* is (Fu, 1995, eq. (18))

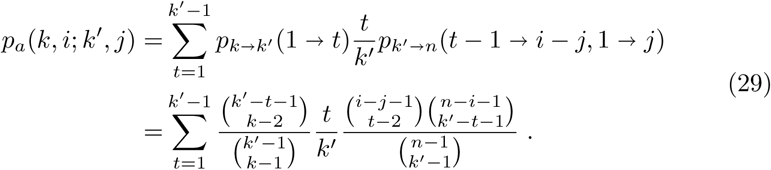

The joint probability that one line at state *k* and one disjoint (not nested) line at state *k^′^* ≥ *k* have *i* resp. *j* descendants at state *n* is (Fu, 1995, eq. (19) and (20))

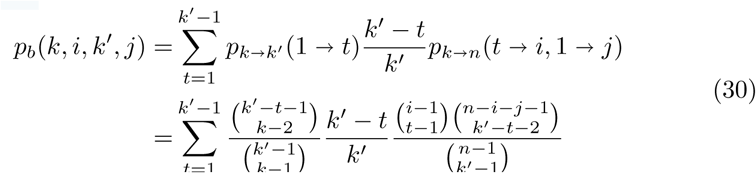

In the latter two cases the summation index *t* runs over the possible numbers of descendants that the line of state *k* may have at state *k′*. Since no single line can be ancestor of all *k′* lines, this number has an upper limit of *k′* – 1. There are more constraints on t as detailled by Fu (1995) (e.g. a line from state *k* can have at most *k′* – *k* + 1 descendants at state *k′*, hence only *t* ≤ *k′* – *k* + 1 is meaningful), however these are already implicit in the binomial coefficients.

Note, that Fu defined these equations only for the case *k′* < *k′*. Using the special definition for the binomial coefficient, they include the case *k′* = *k′* (Durrett, 2008): if the lines are from the same state, then *t* = 1 and we have 
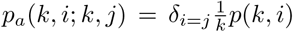
and 
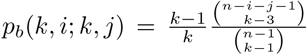
. These two equations correspond to eq. (14) and (15) of (Fu, 1995).

Hence the probability, that a line at *k* and a line at state *k* ≤ *k*′ have *i* respective *j* descendants at state *n* yields for 2 ≤ *k* ≤ *k*′ ≤ *n*:

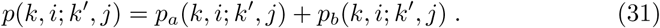

Now we derive the probabilities involving three lines. These may be all of the same state, of two different states or of three different states. We assume *k* ≤ *k′* ≤ *k″*. We take a single line at each state *k*, *k′* and *k″*respectively and subdivide along their possible relationships. We denote the lines *l*, *l″* and *l″* respectively. The six cases are (compare figure 8):

- *aa*: *l′* is a descendant of *l* and *l″* is a descendant of *l′*
- *ab*: *l′* and *l″* are both descendants of *l*, but *l″* is not a descendant of *l′*
- *ba*^(3)^: *l′* is a descendant of *l*, but *l″* is not
- *ba*^(2)^: *l″* is a descendant of *l*, but is not *l′*
- *ba*^(1)^: *l″* is a descendant of *l′*, but both are not descendants of *l*
- *bb*: no line is a descendant of any of the other two lines

As before, *t* counts the number of descendants of line *l* at state *k′*. *t*_1_ denotes the number of descendants of *l* at state *k″*, without the descendants of *l′*. *t_2_* finally counts the descendants of line *l′*. We present here only the first case, while all six cases are listed in the supplement.

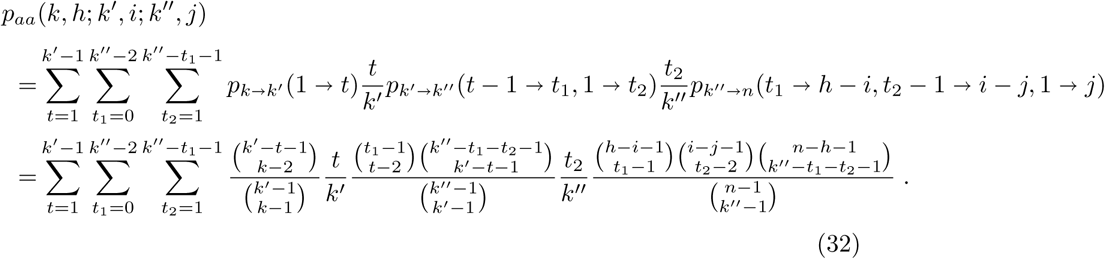

Since the six cases cover all possible combinations, the total probablity that three lines at state *k*, *k′* and *k″* resp. (with *k*, ≤ *k′* ≤ *k″*) have *h*, *i* and *j* resp. descendants at state *n* is given by

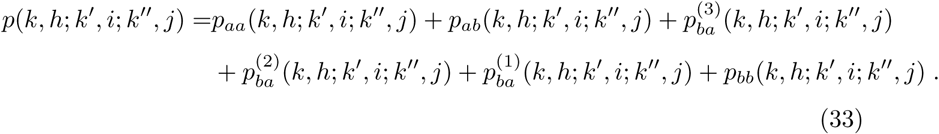

We now relate the indicator variables of eq. (24) to the above probabilities. For two lines we have the three cases distinguished by Fu (1995, text and equations without number, before eq. (22))

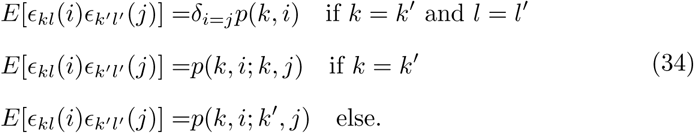

With three lines (still assuming *k* ≤ *k′* ≤ *k″*, this extends to:

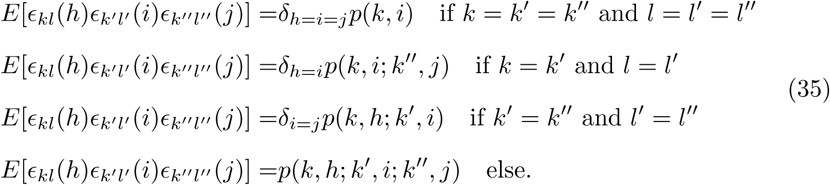

#### 3.1.3. Averaging over line lengths

**Proposition 1.** *For any* 1 ≤ *k*, *k′*, *k″* < *n*, 1 ≤ *l* ≤ *k*, 1 ≤ *l′* ≤ *k′*, 1 ≤ *l″* ≤ *k″ the following equation holds:*

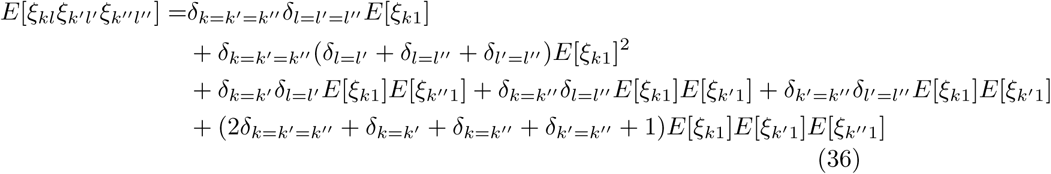

Proof. Let *X* be a random variable. It can be easily shown that, if *X* is exponentially distributed (*X* ~ *Exp*(λ)), then the first three moments of *X* are 
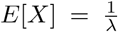
,

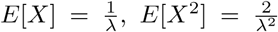
and 
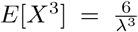
. If *X* is is Poisson-distributed (*X* ~ *Poisson*(*μ*)), than *E*[*X*] = *μ*, *E*[*X*^2^] = *μ* + *μ*^2^ and *E*[*X*^3^] = *μ* + 3*μ*^2^ + *μ*^3^. In agreement with the definition of the coalescent the *ξ_kl_* are distributed as 
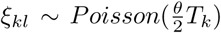
 with 
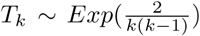
. *ξ_kl_* and *ξ_k′l′_* are independent if *k* ≠ *k′* while *ξ_kl_* and *ξ_kl′_* are independent conditional on *T_k_* for *l* ≠ *l′*. We follow here an analogous derivation as in Wakeley (2008).

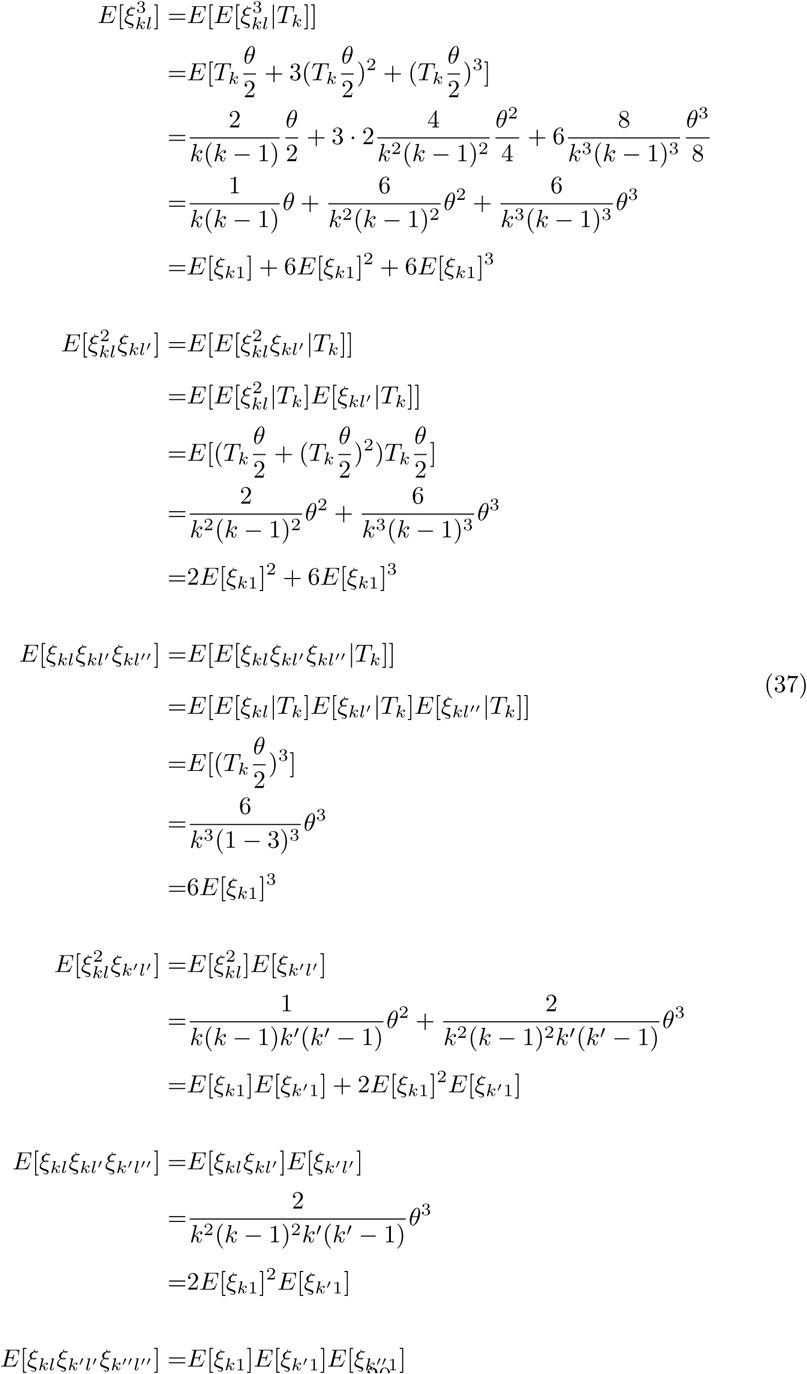

#### 3.1.4. Combining results

We insert now the results for averaged topologies and averaged line lengths into eq. (24):

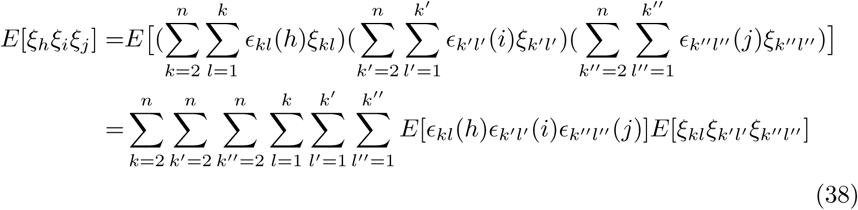

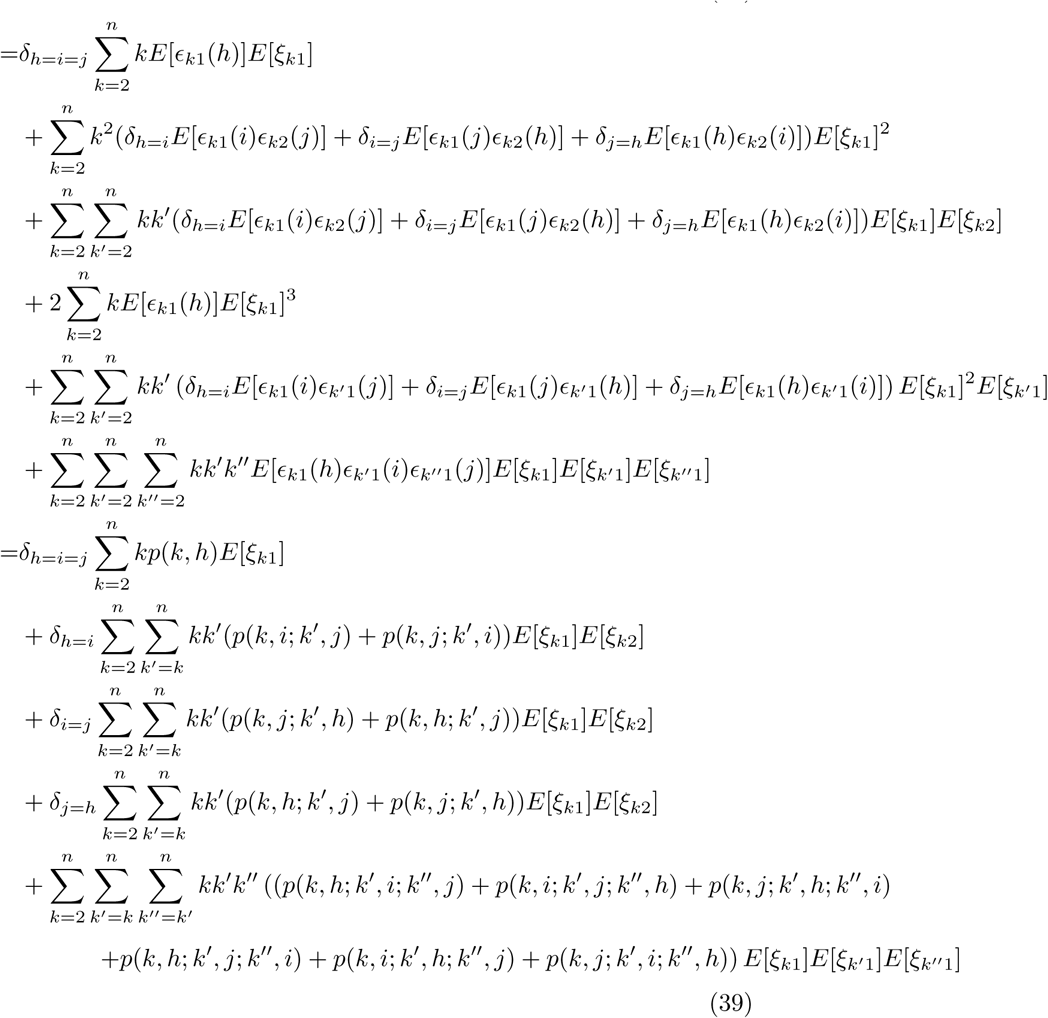

Applying eq. (22) of (Fu, 1995) to the first term of (39) yields eq. (2):

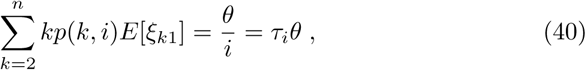

and applying his eq. (23) to the next three terms of (39) yields eq. (4):

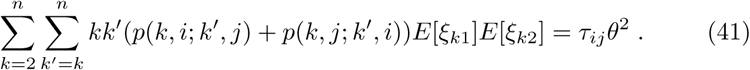

We now define the remaining terms (39) as functions

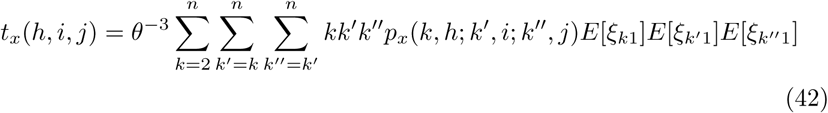

where *x* stands for {*aa*, *ab*, *ba*^(3)^, *ba*^(2)^, *ba*^(1)^, *bb*} and finally we set

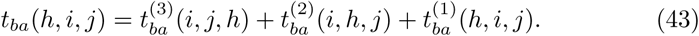

In the supplement we transform these functions to yield (6).

We offer an implementation in C++ for numerical calculation of the third moments, given *n* and *θ*, using the expressions (1)-(6). Just for control, we implemented the unsimpli ed functions (42), too. Within rounding errors (< 10^−12^) they yield the same values as (6) for all third moments *E*[*ξ_h_ξ_i_ξ_j_*] and tested sample sizes 2 ≤ *n* ≤ 17. With the algebraic computing software Mathe-matica (Wolfram Research, Inc., 2014) we were able to prove for the same range of *n* that the expressions are exactly equivalent. The source code is con-tained in the package “coatli”, downloadable at http://sourceforge.net/projects/coatli.

### 3.2. Proof of theorem 2

We derive the third moments of segregating sites *S* using the method of Watterson (1975). He showed (his eq. (1.3a)), that the probability generating function of *S* can be approximated for large population size *N* and small sample size *n* by:

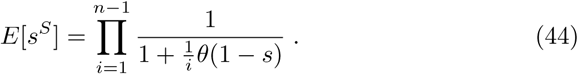

From general probability theory we use the formula

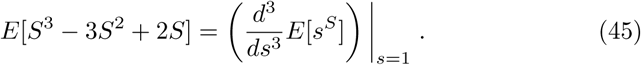

Hence:

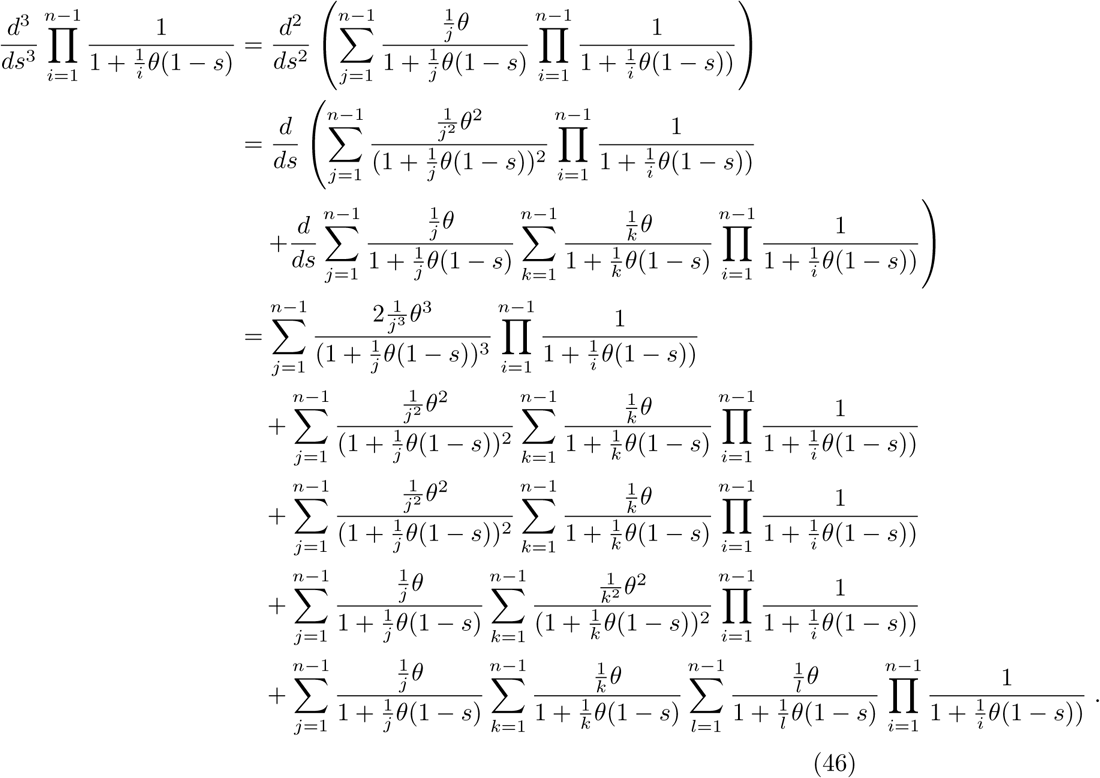

Setting *s* = 1 gives

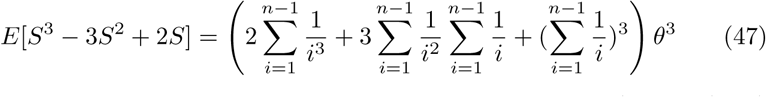

and inserting Wattersons results for the first and second moment (his eq. (1.4a) and (1.5a))

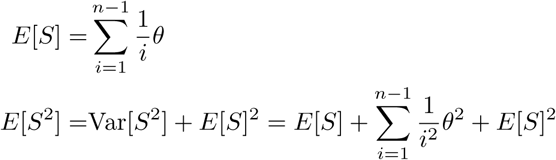

Yields our theorem 2:

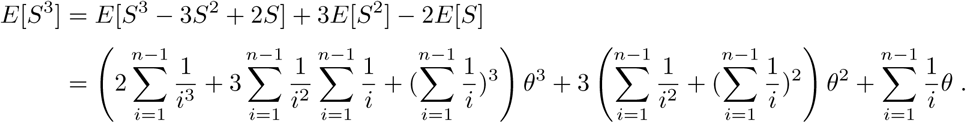

## 4. Discussion

Kingman’s coalescent (Kingman, 1982) is an extremely useful model to describe the patterns of mutations in neutral populations. For this reason, coalescent methods were used to compute analytically the expectation and covariance of the frequency spectrum (Fu, 1995). Here, we derive for the first time the third moments of the full frequency spectrum. We think, the third moments add a valuable building block to coalescent theory.

Beyond their fundamental interest, our results have several applications. We show how to compute analytically the bias of neutrality tests. Moreover, we describe the joint frequency spectrum for triplets of sites (fully characterising their expected haplotype structure). In turn, these results can be used to improve neutrality tests and approaches based on composite likelihood (Kim and Stephan, 2002) and Poisson random field (Sawyer and Hartl, 1992).

The conditional spectra can be used to characterize chromosomal inversions and introgressions (Ferretti et al., 2017). The evolution of inversions has been studied already a long time (Corbett-Detig and Hartl, 2012). Recent improvements of high-throughput sequencing technology allow their investigation on a much larger scale (Sudmant et al., 2015). When alleles are found at intermediate frequency, it is not obvious, whether they are under balancing selection, ongoing positive selection or just neutrally evolving by genetic drift (Hoffmann and Rieseberg, 2008). Patterns of polymorphisms in such regions may help to tackle this question. In regions with inversions, recombination can be strongly inhibited (Kirkpatrick, 2010) which allows to partition the spectrum into nested and disjoint components with respect to the inverted sequences. Nested/disjoint spectra can hence be used to extend the class of frequency spectrum based tests on neutrality to cope with genomic features such as inversions and introgressions. The proper normalisation of such tests requires the knowledge of the corresponding variances and covariances, which we derived.

The main limit of our results is that they do not account for recombination between sites. Recombination is largely irrelevant for the spectrum of a single site, but becomes already relevant for pairs of sites. Therefore, applicability to biological data is limited to small regions or sequences with negligible recombination. For this reason we present an application to the dynamics of chromosomal inversions. The patterns of mutations in these regions are naturally described in terms of the higher moments of the frequency spectrum without recombination. In particular, the expected spectrum of neutral inversions can be obtained from the second moments of the usual spectrum, as shown in Ferretti et al. (2017), while the variance of the spectrum of neutral inversions requires precisely the third moments of the spectrum derived here. Applications to the detection of balancing and positive selection in chromosomal inversions and other structural variants will be presented in future publications.

Note that there is a close relation between the joint spectrum of multiple sites and the multi-allelic spectrum of a single locus (Ferretti et al., 2017). In fact, at low mutation rates, we can consider the multiple sites as a single locus with multiple alleles, and retrieve the multi-allelic spectrum for the locus by considering the frequencies of the *m* + 1 alleles that result from the *m* polymorphic sites. In this light, our results can be used to derive the full quadri-allelic frequency spectrum. This could be applied to several multiallelic variants, the more relevant being nucleotide polymorphism (which have at most four alleles A,C,G,T). Related results can be found in Jenkins and Song (2011) and Bhaskar et al. (2012).

The results in this paper apply to a sample of size *n* much smaller than the size of the population. The spectrum for large samples converges to the continuous population spectrum for triplets of sites. It would be interesting to derive analytically simple expressions for such a spectrum, similarly to Ferretti et al. (2017). However, our expressions contain many explicit sums that prevent a direct computation. A further simplification of the expressions provided in this paper would be helpful.

## Acknowledgment

We thank Iulia Dahmer and Götz Kersting for insightful discussions. AK was supported by a grant of the German Science Foundation (DFG-SFB680) to T. Wiehe (University of Cologne).

## Footnotes

1 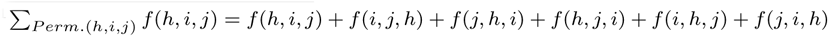

2 From the general expansion (e.g. Van Erp and Van Gelder (2007))) 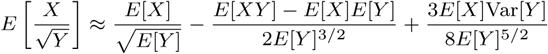

